# Experimentally quantifying impact of herbivory on duckweed communities in natural pond ecosystems

**DOI:** 10.1101/2023.08.08.552541

**Authors:** Swapna K. Subramanian, Martin M. Turcotte

## Abstract

Plant herbivory structures communities, impacts energy and nutrient flow in ecosystems, and drives speciation. Yet, our knowledge of plant-herbivore interactions remains limited in freshwater ecosystems compared to terrestrial ones. We still lack robust experimental data on the impact of herbivores in nature on whole families of important macrophytes such as the globally distributed duckweed family (Lemnaceae). We conducted a replicated manipulative field experiment, using exclosures, across multiple bodies of water quantifying both ambient herbivory as well as herbivory caused by the addition of two different weevil and aphid herbivores. We found that invertebrate herbivores can strongly impact duckweed multi-generational population growth (e.g., reducing daily relative growth rate by up to 82% compared to controls) and differentially impact duckweed species composition. These impacts, however, vary greatly across sites and with the identity of the herbivores. Our results suggest that insect herbivores can severely slow the growth of the of the world’s fastest growing plant family. It also provides crucial information as duckweed continue to be developed for various applied purposes such a biofuel production and bioremediation.

## Introduction

The consumption of plants by herbivores is a critical biotic interaction structuring natural and human dominated ecosystems (McNaughton et al. 1989; Huntly 1991; Coley and Barone 1996; Poore et al. 2012; Bakker et al. 2016). Herbivores can strongly impact plant population dynamics, alter species composition, energy and nutrient flow through ecosystems, and drive the evolution of defense and cause speciation (Cyr and Pace 1993; Gruner et al. 2008; Schmitz 2008; Turcotte et al. 2014a; Agrawal and Zhang 2021). Yet, for many decades most of the research on plant-herbivore interactions were primarily conducted in terrestrial ecosystems, and the prevailing view was that herbivory of aquatic plants was not important. Lodge’s (1991) as well as Cyr and Pace (1993) seminal reviews of the literature had a critical impact in not only amassing evidence of the importance of consumers on macrophytes but also galvanized more research on this topic (Bakker et al. 2016). Although we now have much more evidence for their importance, our understanding of macrophyte-herbivore interactions remains limited especially in freshwater systems (reviewed in: Poore et al. 2012; Bakker et al. 2016; Wood et al. 2017; Pulzatto et al. 2018).

Herbivory can significantly impact aquatic macrophytes resulting in strong community and ecosystem effects (Jacobsen and Sand-Jensen 1992; Bolser and Hay 1998; Bakker et al. 2016; Wood et al. 2017). Although herbivores can strongly impact macrophyte populations and communities, their effect sizes can vary in space and time because of heterogeneous abiotic and biotic factors including resource availability and herbivore community composition (Huntly 1991; Lodge 1991; Tomas et al. 2005; Wood et al. 2017). In fact, Bakker et al. (2016) suggested that herbivory was much more variable, and on average 5-10 times higher, in aquatic ecosystems than reported in Turcotte et al.’s (2014a, b) review of terrestrial herbivory. Numerous hypotheses based on ecosystem properties, and various aspects of both the plant and herbivore species involved such as stoichiometry, growth rates, and growth form, have been hypothesized to explain such differences (Cyr and Pace 1993; Cebrian 1999; Prado et al. 2010; Bakker et al. 2016). One factor that clearly matters across ecosystems is the identity of the consumers and their densities. Wood et al.’s (2017) meta-analysis found that taxonomic groups differed in their impact with insects causing approximately a 28% reduction in macrophyte abundance while vertebrates caused greater reductions on average (44% to approximately 58%). Moreover, abiotic factors including the availability of resources such as light and nutrients may also drive variability in herbivory (Coetzee et al. 2007; Gruner et al. 2008; Hidding et al. 2016; Bakker et al. 2016). Herbivory and abiotic conditions can also interact to influence plant species composition (Coley and Barone 1996). For example, Center et al. (2005) showed that specialist herbivores and nutrient availability alter the competitive dynamics between two floating aquatic macrophytes. In summary, although our understanding of herbivore-macrophyte interactions is growing in numerous study systems, it remains surprisingly limited in others.

The duckweed family (Lemnaceae), sometimes referred as the subfamily Lemnoideae within the Araceae, are a globally distributed group of 37 species of floating macrophytes that can dominate freshwater ponds (Landolt 1986; Tippery and Les 2020) with significant ecosystem level impacts (Rabaey and Cotner 2022). Given their rapid reproduction of only a few days, when reproducing asexually, they have become an emergent model system to study evolutionary-ecology and genomics of species interactions, environment-host interactions, plant-development, and adaptation (Laird and Barks 2018; Sandler et al. 2020; Tan et al. 2021; Acosta et al. 2021; Hess et al. 2022). They are also being developed as a biofuel, a bioremediation agent, a forage crop, and to produce drugs (Appenroth et al. 2015; Ekperusi et al. 2019; Acosta et al. 2021; O’Brien et al. 2022). Even though this group of highly simplified formerly terrestrial plants has been the focus of study for many decades (Guppy 1894; Hillman 1961; Landolt 1986), studies of interactions among duckweed and their herbivores remain nascent.

Beyond surveys of herbivore communities on duckweed, experiments remain very rare. Laboratory experiments usually focus on herbivore preference (Mansor and Buckingham 1989) and/or quantify the impact of herbivory on different duckweed species or genotypes (Pípalová 2003; Lee et al. 2022; Schäfer and Xu 2022). For instance, Subramanian and Turcotte (2020) showed that the water-lily aphid (*Rhopalosiphum nymphaeae*) prefers specific duckweed species and these species have differential tolerance and resistance to herbivory which might adhere to the resource availability hypothesis (Endara and Coley 2011). Mariani et al. (2020), found that a native moth could consume and strongly harm an invasive duckweed species in Italy. Outdoor mesocosm experiments are rarer and are more challenging than lab experiments because they allow for more confounding environmental factors, e.g., weather, while still being more controlled than true field experiments. For instance, Tipping et al. (2009) manipulated the presence and frequencies of a duckweed and an aquatic fern, as well as nutrients, and the presence of a weevil that could only feed on the fern. They found complex interactions among these factors but no indirect effect of herbivory impacting the duckweed.

Although more difficult to conduct, field experiments provide a more realistic approach to quantify the impact of herbivores on macrophytes (Paine 1980; Pulzatto et al. 2018). Unlike, terrestrial and marine systems wherein field experiments consisting of exclosures, e.g., deer fences, are very common (Poore et al. 2012), we know of only a single study doing so in duckweed. Carlsson and Lacoursière (2005) tested the impact of an invasive snail using exclosures in one wetland. Adding one snail, or none, they found an almost complete eradication of the duckweed within only 6 days. Although showing the importance of herbivory, it was only conducted at a single site which limits insight into spatial variation (Lodge 1991). Second, it lacks an ambient herbivory treatment and thus the impact of a single individual was tested. The reason for the rarity could be that such experiments are especially challenging with duckweed because they float implying that the enclosures must keep them contained while allowing access by the herbivore community that may be swimming, flying, or walking on the water surface. Thus, whether duckweed with their immensely rapid rate of asexual reproduction (Ziegler et al. 2015) are impacted by herbivores in nature remains unresolved.

To fill this gap, we conducted a field exclosure experiment focusing on invertebrate herbivores replicated within and across four ponds. We quantified the multigenerational performance of a mixture of two duckweed species growing without herbivores, exposed to ambient herbivory, and to either aphids or weevils. In doing so we addressed: Q1) Are duckweed communities significantly harmed by herbivores? Q2) Do herbivores impact duckweed community composition? Q3) Does the identity of the herbivore matter? Q4) Do these patterns vary across sites?

## Materials and Methods

### Study System

This experiment focused on two abundant duckweed species the Greater Duckweed *Spirodela polyrhiza* (L.) Schleid and the Common Duckweed *Lemna minor* L., both of which are distributed widely across the globe (Armitage and Jones 2020; Tippery and Les 2020). We collected them from Pymatuning State Park, in northwestern Pennsylvania, USA in 2017. We established laboratory colonies of each species from single individuals that only reproduced clonally under laboratory conditions. The experiment used a single genotype per species to avoid rapid evolutionary dynamics (Hart et al. 2019; Malacrinò et al. 2022) that could potentially confound our results. Colonies were maintained at room temperature on windowsills free of any herbivores for a week preceding the experiment and maintained on 50% concentration growth media (Appenroth et al. 1996).

Similarly, in 2017 we established a laboratory colony of water-lily aphids from a single individual collected from Twin Lakes Park Westmoreland County, Pennsylvania, USA. The aphid was collected form a community of duckweed including *S. polyrhiza, L. minor,* and *Wolffia brasiliensis*. The aphids were maintained on colonies of *S. polyrhiza* and *L. minor* growing at room temperature on windowsills which keeps them reproducing asexually. For two weeks preceding the experiment they were grown on *S. polyrhiza* from Twin Lakes Park. Natal host has limited impact on aphid preference (Subramanian and Turcotte 2020). In addition, we used the widely distributed Duckweed Weevil (*Tanysphyrus lemnae*) which is considered one of the most common and widespread duckweed herbivores, but, to our knowledge, no study has shown the impact of this insect on duckweed population dynamics (Center et al. 2002; Lee et al. 2022). Weevils were collected the day preceding the experimental setup from the site Shelter Nine, Pymatuning State Park. Weevils were kept on *S. polyrhiza* and *L. minor* collected from this site until placement on experimental duckweed.

### Experimental Procedure

We selected four still bodies of water within the Pymatuning State Park (Fig. 1), that all had almost total duckweed cover but did vary in duckweed species composition. The four sites are Shelter Nine a pond located a short distance from Pymatuning reservoir (41.506991, - 80.457792), Spillway Close an inlet off Pymatuning Reservoir (41.630392, -80.441723), Spillway Far a pond near Pymatuning Reservoir (41.632432, -80.439675), and Stuart’s Bay a small bay off Pymatuning Reservoir (41.645797, -80.431482). All sites had *L. minor* and *S. polyrhiza* in varying amounts (Fig. 1), and Stuart’s Bay had a small quantity of *Wolffia brasiliensis*, a tiny species of duckweed that aphids cannot consume to our knowledge (Subramanian and Turcotte 2020). We constructed floating chambers to manipulate duckweed herbivory (Fig. 2). Each consisting of clear, plastic, one-gallon square wide mouth containers (dimensions of 260 (h) × 145 × 128 mm, code #0070-08, SKS Bottle and Packaging, USA), with white lids. Using a hot knife, 15×8 cm openings were cut out from three sides. A 7.5 × 7.5 cm square was cut out from the plastic lid. We glued various meshes over the openings to manipulate herbivore treatments using Gorilla Glue ® All-Temperature hot glue (Joann Fabrics, USA). White casa chiffon fabric (code #15079494, Joann Fabrics) was used as our fine mesh with holes less than 1 mm, and white petticoat netting fabric (code #1825165, Joann Fabrics) with holes approximately 6.35 mm as our more open mesh. We glued fine mesh onto all lids regardless of herbivore treatment. Four such chambers were attached together and to a 13×13 cm piece of polystyrene foam with a hole in the center (code #304090, Lowe’s, USA) using plastic threaded rods, hex nuts, and washers (U.S. Plastic Corp, USA). We then hammered a garden stake into the pond sediment and slid the apparatus onto the rod preventing horizontal movement but allowing vertical movement as water depth changes (Fig. 2). Each apparatus was replicated four times in each site (placed at least 2 m apart and usually farther).

**Figure 1:**
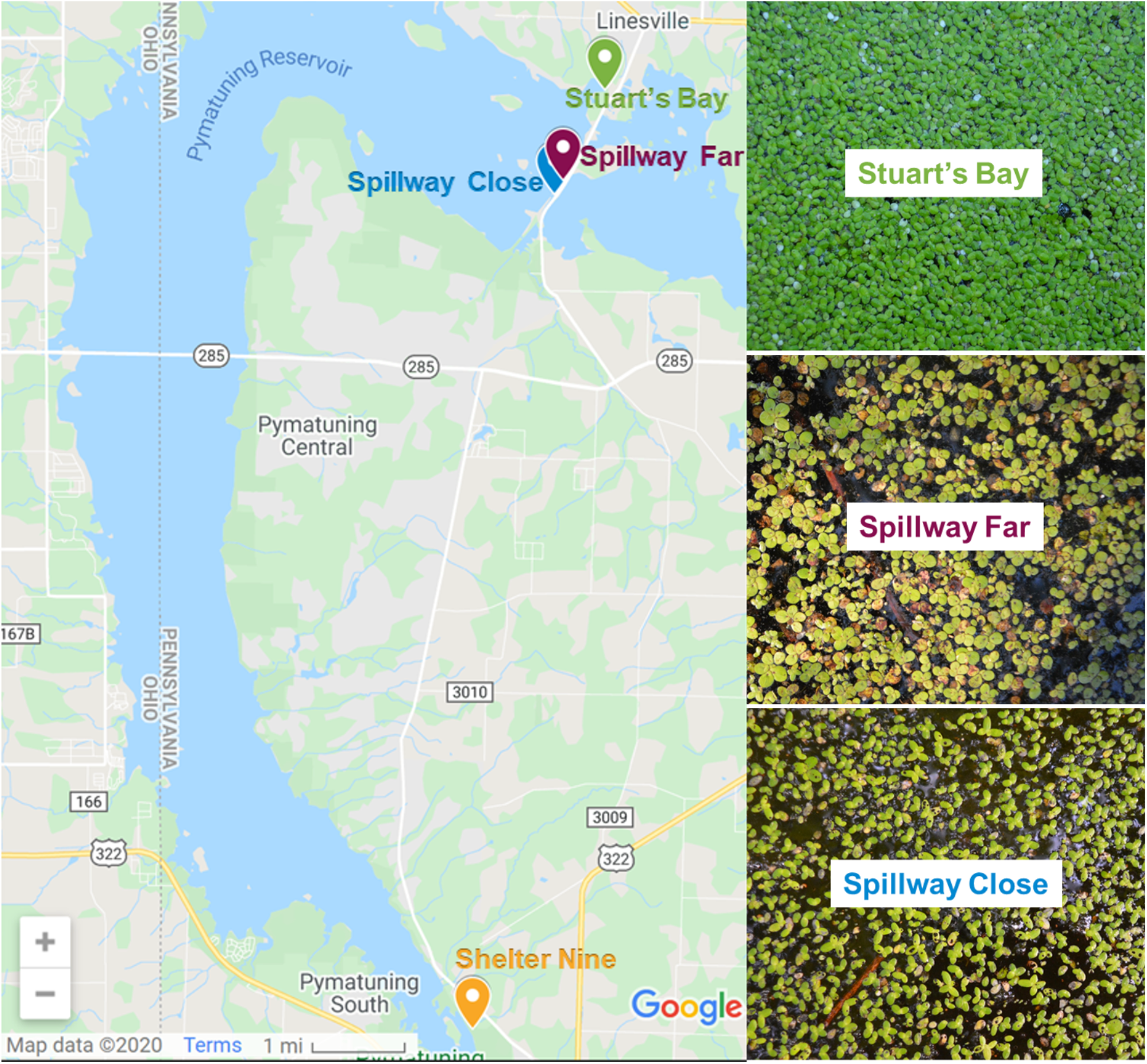
Locations of field experiments and representative image of natural species composition at each site in Pymatuning State Park, Pennsylvania, USA. No photo of Shelter Nine is presented because of a complete loss of this site due to a drying event.

**Figure 2:**
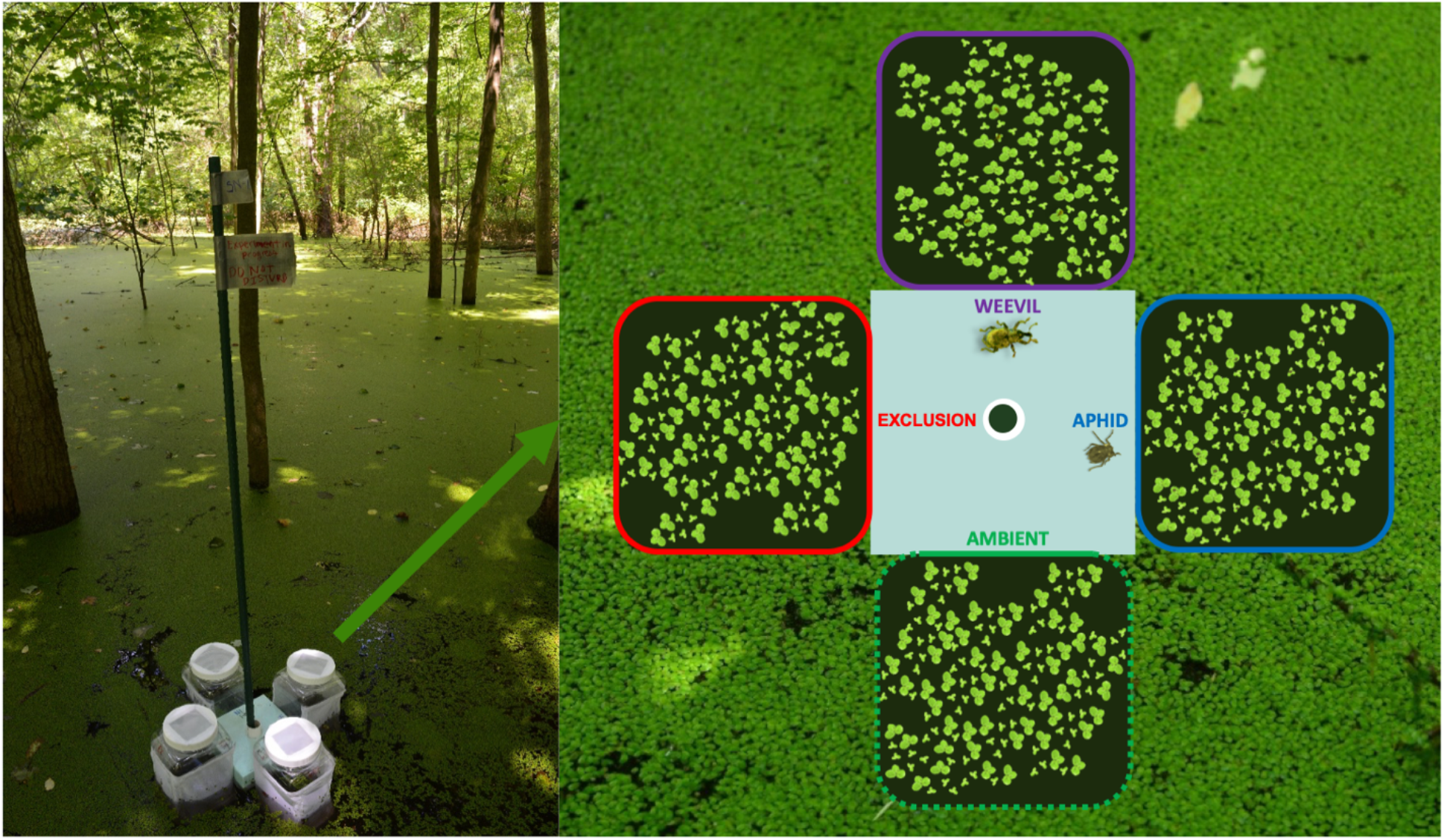
Floating field experimental chambers to contain duckweed and manipulate herbivore composition using mesh. Photos are from field site Shelter Nine.

On July 2^nd,^ 2019, in each replicated chamber, we added 60 individuals of *S. polyrhiza* and 90 of *L. minor* with approximately equal surface areas. In each quad-chamber apparatus we established the following herbivore treatments: herbivore exclusion, ambient herbivory, and addition of either the aphid or the weevil only. The ambient herbivory treatment, which quantifies natural rates of herbivory from small herbivores such as insects and snails, used the open mesh that had wide enough holes (∼6.35 mm) to allow most aquatic, walking, and flying invertebrate herbivores entry. The herbivore exclusion treatment used a very fine mesh that excluded all herbivores in the field. The addition treatments also used the fine mesh. In the aphid treatment we added 15 3^rd^ instar aphids and the weevil treatment received 5 adult weevils. The quantity of herbivores added were similar to those observed on equal areas of duckweed as the chamber (∼156 cm^2^) in various locations (S. Subramanian, *personal observation*).

We collected additional data to provide more insight into spatial variation in our results. We surveyed herbivore composition in the site and within the ambient herbivory treatment but given logistical constraints we could not acquire quantitative estimates. Herbivore samples were collected from each site during the experiment as well as from the ambient herbivory treatment at the end of the experiment and preserved in 70% ethanol. In addition, on the last day of the experiment, water samples were taken from below each experimental apparatus and concentrations of Nitrate as well as Total Phosphorus were quantified by the Pennsylvania State University Agricultural Analytical Services Laboratory.

### Data Collection and Analysis

To quantify community impacts of the various forms of herbivory we measured community size, as the total area and abundance of each duckweed community at the end of the experiment. Sites differed in how long the experiment lasted, between 21 and 24 days, and thus we calculated and analyzed relative growth rate (RGR) as:

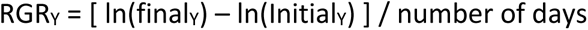

where Y is the parameter area or abundance (N). RGR is this context is equivalent to daily exponential growth rate (Ziegler et al. 2015). We ran models on two different datasets, one with the exclusion and ambient herbivory treatments only, and the other dataset that included the exclusion and herbivore addition treatments (aphid addition and weevil addition) only. Herbivore impact on community RGR_Area_ and RGR_N_ were separately analyzed using linear models with herbivore treatment and site as fixed effects and tested factor significance using ANOVA. In addition, we tested herbivore impact on species composition by calculating the proportion of area represented by *S. polyrhiza.* We used generalized linear models with site and herbivore treatment as fixed effects. We then analyzed these results using a *chi*-squared test ANOVA. Effect sizes for both types of data were based on estimated marginal means and calculated as percent change from the exclusion treatment (herbivore treatment-exclusion treatment)/exclusion treatment.

## Results

Shelter Nine was lost due to a severe drying event that grounded all chambers killing the duckweed. Water chemistry analysis of each site revealed that all three had nitrate levels below the detection limit (<0.02 mg/L). For total phosphorous, Stuart’s Bay had an order of magnitude higher concentration (0.26 ± 0.008 mg/L) than Spillway Close (0.035 ± 0.004 mg/L) and Spillway Far was below the detection limit (<0.025 mg/L).

### Ambient Herbivory

Ambient herbivory treatments worked as intended and the mesh allowed both crawling and flying aquatic invertebrate duckweed herbivores in to consume the duckweed. We found that herbivore composition varied between sites. Through observations of damage type and herbivore presence in ambient herbivory treatments, we determined that all three sites had Duckweed weevils (*T. lemnae*), Duckweed flies (*Lemnaphila scotlandae),* and Springtails (*Podura aquatica*). However, Stuart’s Bay also had Water-lily aphids (*R. nymphaeae*) and Spillway Far had the Duckweed moths (*Elophila spp.*).

Our experiment revealed that ambient herbivory reduced duckweed relative growth rate and varied among sites. The impact of ambient herbivore differed greatly among sites (Fig. 3, marginally significant interactions for RGR_Area_ *p* = 0.08 and RGR_N_ *p* = 0.07). For instance, Spillway Far being much more heavily damaged (-82% loss of RGR_Area_ and -69% loss of RGR_N_) and other sites much less. Averaging across the sites, ambient herbivory reduced RGR_Area_ by - 42% (ANOVA, *p* < 0.001) and decreased RGR_N_ by -31% (*p* = 0.001). Site was also highly significant (both *p < 0.002)* with slower growth in Spillway Far (Fig. 3).

**Figure 3:**
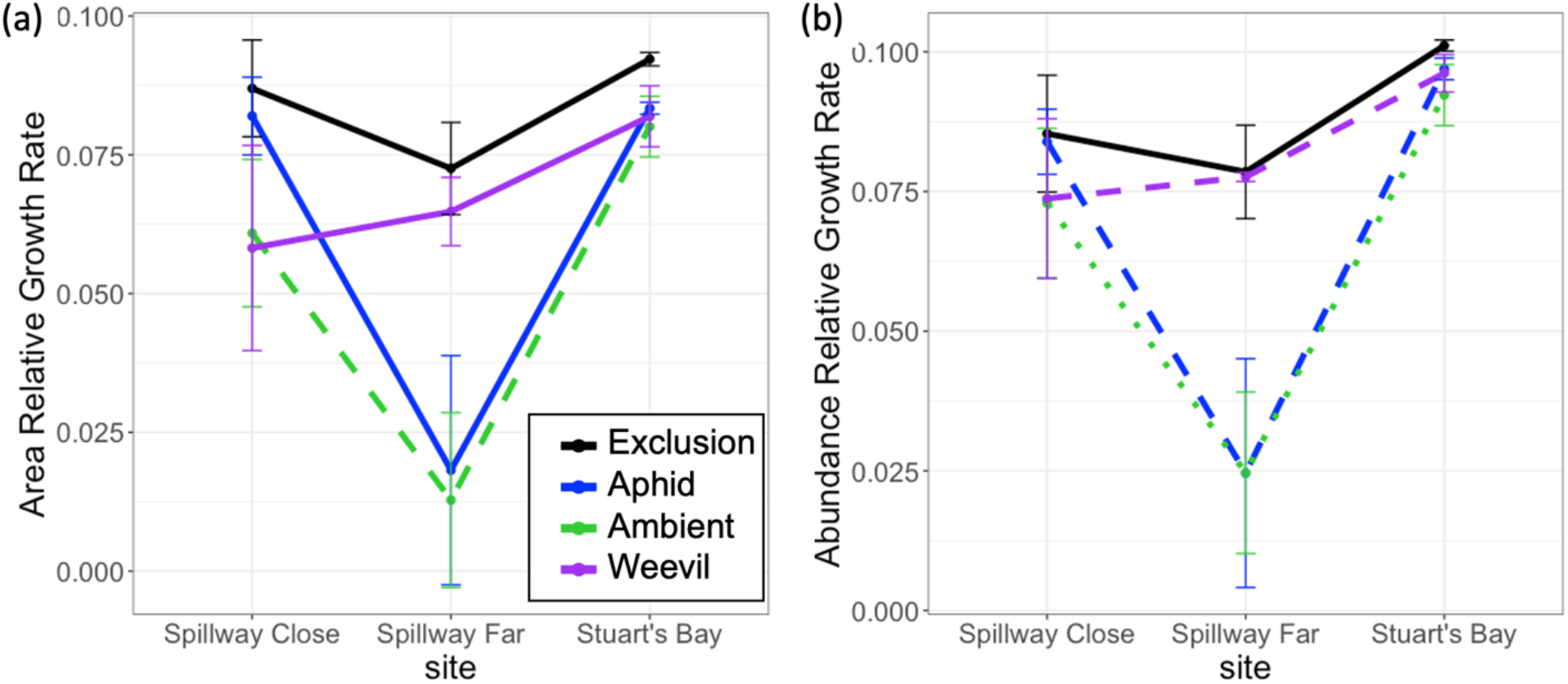
Relative Growth Rate (RGR) of duckweed community (a) area and (b) abundance across the three surviving sites and the four herbivory treatments. Errors bars represent ±1 S.E.

Duckweed species composition was significantly impacted by an interaction between herbivory and site (Fig. 4, χ^2^, interaction *p*< 0.001) but not main effects (*p > 0.35*). Ambient herbivory decreased the proportion of *S. polyrhiza* at Spillway Close (-27%) and Stuart’s Bay (- 20%) but a large increase of +106% in Spillway Far compared to site control treatments.

**Figure 4:**
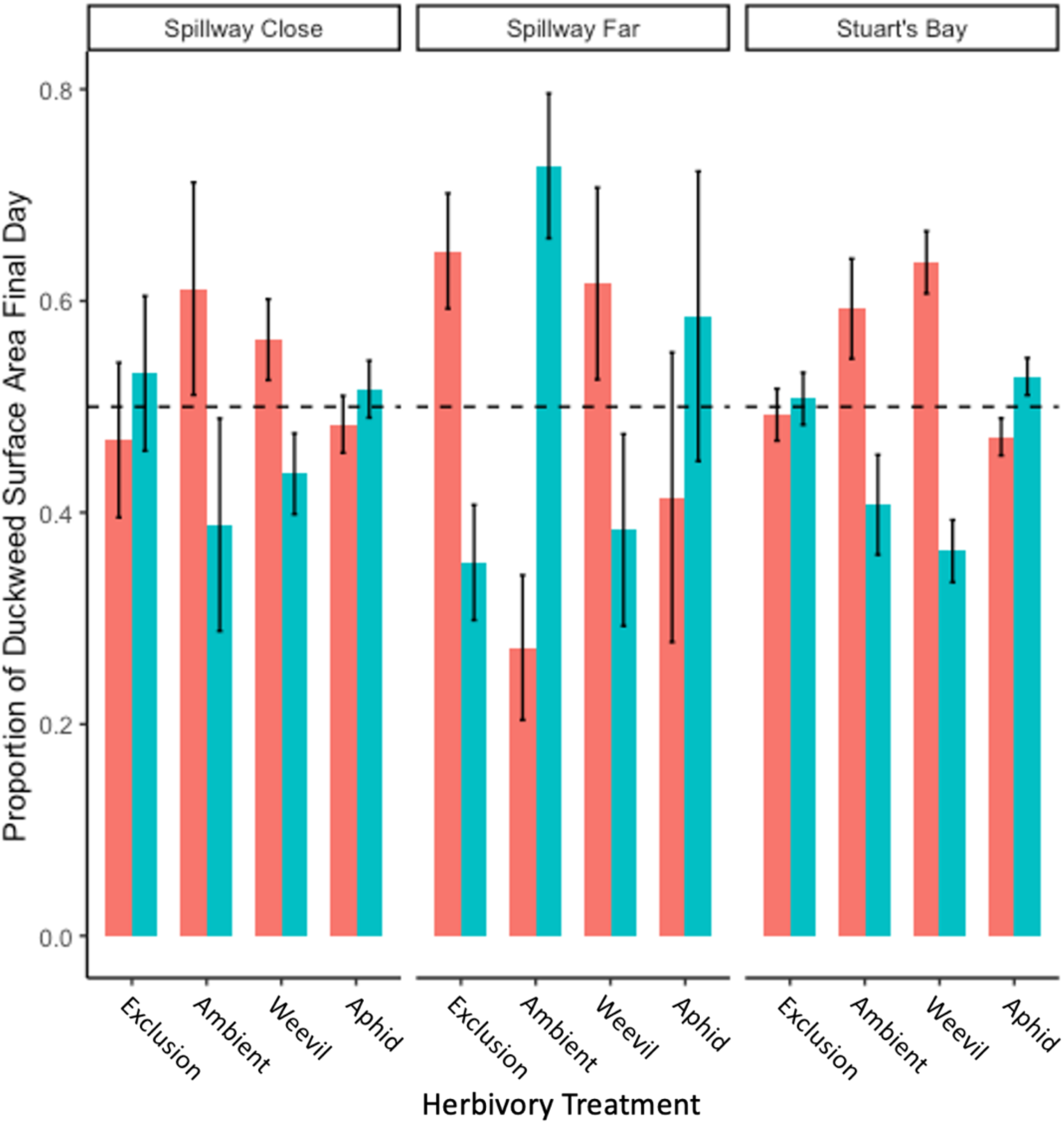
Final proportional duckweed area by species across sites and the four herbivory treatments. *Lemna minor* is in red and *S. polyrhiza* in teal. Errors bars represent ±1 S.E.

### Herbivore Addition Treatments

Adding either weevils or aphids reduced the growth rate of the duckweed community, but this impact interacted with site (Fig. 3). Analyzing the exclusion, weevil, and aphid treatments together revealed that RGR_Area_ was influenced by all factors (ANOVA, herbivore treatment: *p*= 0.00389, site: *p*= 0.0031, interaction: *p*=0.0470) and so was RGR_N_ (herbivore treatment *p*= 0.00364, site: *p* = 0.0007, interaction: *p*= 0.0317). Duckweed species composition however was only influenced by herbivore treatment (χ^2^, *p*= 0.0257, Fig. 4).

Specifically looking at the impact of aphids compared to the exclusion treatment aphids reduced RGR_Area_ by -30% and RGR_N_ by -25% on average (Fig. 3). Yet clearly, there is large spatial variation as their impact ranged from -6% to -75% reduction in RGR_Area_. Aphid addition had little impact on the relative performance of the duckweed except in Spillway Far where it caused an increase of +66% in *S. polyrhiza* final area (Fig. 4) compared to the exclusion.

Weevil addition overall reduced the total duckweed RGR_Area_ by on average -18% but also and varied about 3-fold across sites. Weevils had a much smaller impact on RGR_N_ reducing it on average by only -7% but this also varied among sites from -1% to -14%. *Spirodela polyrhiza* generally performed worse or equally well when exposed to our weevil treatment than in the control with a mean decline in percent final area of -12% but this varied among sites.

## Discussion

Only in recent decades has the importance of herbivory on freshwater plants been demonstrated in contrast to the long-standing tradition in terrestrial systems (Lodge 1991; Bakker et al. 2016; Wood et al. 2017). Here we report on the second ever experimental study quantifying herbivory in duckweed in nature even though this is a globally distributed family of plants of growing importance in basic research and for biofuel and bioremediation (Appenroth et al. 2015; Laird and Barks 2018). Our replicated manipulative field experiments across multiple bodies of water, using three different herbivore treatments, showed that invertebrate herbivores can strongly impact duckweed multi-generational population growth and species composition. Yet, these effects strongly vary both quantitatively and qualitatively among sites and with the identity of the herbivore.

### Community Size

Even though duckweed may be the world’s fastest growing plant (Ziegler et al. 2015), insect herbivores can significantly impact their growth rates. We found that ambient insect herbivory caused strong reductions in population growth rate of the duckweed community but in a context dependent manner. Daily exponential growth rates calculated as area or abundance of were reduced by 42% and 31% respectively across our three sites. These are very strong negative impacts on growth rates and if we calculate missing area using these RGR values, after 24 days it would represent a loss of 54% in area and 45% in abundance. This is a large impact especially if we consider that we excluded any herbivores larger than our mesh holes (6.35 mm). These are much higher rates of damage than the average yearly loss of 5.3% leaf area estimated in terrestrial plants (Turcotte et al. 2014b, a). It is more compared to Wood et al.’s (2017) review of aquatic herbivory showing mean reductions in plant abundances of approximately 28% caused by insects.

We also saw striking quantitative variation among sites with Spillway Far showing much stronger herbivory (Fig. 3). We could not logistically quantitatively track herbivore composition, but we suspect that this is caused by the duckweed moths which was only found in this site. Much of the damage to *L. minor* was chewing and not piercing or leaf mine which suggests the weevils were not as important. Such moths can consume large amounts of duckweed and uses them to create cases during its larval stage (Mariani et al. 2021) and can quickly decimate duckweed populations in this region (M. Turcotte, *personal observation*). A similar species of duckweed moth has been shown to consume an average of 92.5% of the *L. minor* when placed in a petri dish and have been proposed as a biocontrol agent of the invasive species *L. minuta* (Mariani et al. 2020). Overall, our results suggest that the insect herbivore community can greatly impact the fast growth rate of duckweed communities, which could be even more dominant in their absence.

Our insect addition treatments also revealed that a small initial number of aphids and weevils can slow duckweed growth. The 5 adult weevils and the larvae they produced caused the least damage of all treatments, but still a substantial reduction of 18%, their impact was lower for RGR_N_, and varied the least among sites (Fig. 3). We likely did not have any adult recruitment during the experiment as their life-cycle takes about one month (Lee et al. 2022). A longer experiment may have generated a stronger impact of this herbivore. The rapidly reproducing aphids had a stronger impact, although we did initiate the experiment with more individuals. In a 32-day lab experiment Subramanian and Turcotte (2020) found that aphids reduced duckweed abundance by up to 50% compared to controls. We saw even stronger impacts but only in Spillway Far (-82% in RGR_Area_ and -68% in RGR_N_) similar to the ambient treatment. Aphids can asexually reproduced, at similar rates as duckweed, allowing them the potentially regulate fast growing macrophytes (Storey 2007; Subramanian and Turcotte 2020). The strong reduction of duckweed growth rate in Spillway Far is thus not only due to ambient herbivores which were excluded from aphid addition treatments.

It is possible that the lower phosphorous concentration at this site, which is often a limiting nutrient in freshwater systems (Correll 1999), synergized with aphid and ambient herbivory presence to further reduce duckweed growth rate. Duckweed without herbivores grew on average 19% slower in RGR_Area_ at this site than the other two (Fig. 3). Variation in abiotic conditions is thought to impact variation in macrophyte-herbivore interactions (Coetzee et al. 2007; Gruner et al. 2008; Hidding et al. 2016; Bakker et al. 2016). Resource availability can influence tolerance and compensation for herbivore damage, with those plants in higher resources generally having more ability to compensate (Huntly 1991; Hawkes and Sullivan 2001; Center et al. 2005). Moreover, other aspect of the environment can modify plant-herbivore interactions, for instance Van der Heide (2006) found that duckweed consumption by duckweed moths is temperature depended.

### Impacts on Species Composition

We found that herbivory can impact the relative performance of duckweed species even changing dominance, but these effects vary with the ecological context. Stuart’s Bay and Spillway Close had similar patterns in that without herbivore both duckweed species preformed equally well, that aphids had little impact, and that ambient herbivory and weevils harmed *S. polyrhiza* more than the smaller *L. minor* (Fig. 4). These results suggest that *S. polyrhiza* is preferred by various herbivores or is less tolerant of their damage. We do not have information on preference or performance for most of these herbivores except for the water-lily aphid. A series of lab experiments showed that these aphids prefer *S. polyrhiza* over *L. minor* but on the other hand *S. polyrhiza* is more tolerant of damage (Subramanian and Turcotte 2020). It is possible that for this aphid herbivore, at least at these 2 sites, preference disadvantage and tolerance benefit may cancel out.

In Spillway Far, results are very different, here without herbivores *L. minor* is dominant, weevils have little impact, and both ambient and aphids damage *L. minor* much more than *S. polyrhiza.* These qualitative changes in the performance of the species due to herbivory are in sharp contrast to the other bodies of water. One possible mechanism for ambient is a strong preference of duckweed moths for *L. minor,* that are not present at the other sites. Duckweed moth preference remains untested explicitly to our knowledge across these genera. Moreover, *L. minor* is less well defended in terms of phenolic content and resistance to aphids measured as aphid exponential growth rate than *S. polyrhiza* (Smolders et al. 2000; Subramanian and Turcotte 2020). If *L. minor* can grow better than *S. polyrhiza,* potentially due to an ability to maintain growth at lower phosphorus concentrations (but evidence for this idea is mixed see Docauer 1983; Lüönd 1986), giving it an advantage without herbivores but then being very susceptible to the moths in this site. Consistent with this idea is that this pond had very few *L. minor* compared to the other sites (see images in Fig. 1). Why *S. polyrhiza* does so well in this site with the aphids remains unclear. We hypothesize that *L. minor’s* tolerance of aphid damage may be more sensitive to low nutrients, or some other aspect of the site, than *S. polyrhiza*. Clearly more research is needed to explore not only herbivore preference and impacts but also how they interact with environmental conditions (Lodge 1991; Wood et al. 2017).

### Conclusions

This study provides insight into the role of herbivory on plant community productivity and composition in freshwater ecosystems. Studying these effects requires testing multiple sites to reveal interactions with other biotic and abiotic conditions that may quantitatively and qualitatively alter the impacts of various herbivores. Given this complexity, we need more experimental studies in natural communities and hopefully our inexpensive design for enclosures will motivate others to continue testing the impact of herbivory on floating macrophytes. Modifications to the design could be made to test the impact of additional herbivores such as snails which have recently been shown to cause rapid evolution in duckweed (Malacrinò et al. 2022). Addressing the dearth of studies on duckweed-herbivory is critically important as we further reveal their ecological importance (de Tezanos Pinto et al. 2007; Rabaey and Cotner 2022), try to control duckweed species that have become invasive (Reeves and Lorch 2012; Mariani et al. 2020), and their expanding use for large scale bioremediation and biofuel production (Appenroth et al. 2015) .

## Acknowledgements

We thank the Department of Conservation and Natural Resources for permission to conduct the experiments in Pymatuning State Park. We thank Julie Everett and Joshua Armstrong for their help conducting the experiment. S.K.S. was supported by a University of Pittsburgh Pymatuning Lab of Ecology Leasure K. Darbaker Prize in Botany and M.M.T. was supported by an NSF grant (#1935410).

## Author Contributions

SKS and MMT conceived the research and designed the experiment. SKS performed it and both authors analyzed the data and wrote the manuscript.

